# Effects of annual rainfall on dengue incidence in the Indian state of Rajasthan

**DOI:** 10.1101/423517

**Authors:** Nasir Salam

## Abstract

Dengue has become a major public health problem in the last few decades with India contributing significantly to the overall disease burden. Most of the cases of Dengue from India are reported during Monsoon season. The vector population of dengue is affected by seasonal rainfall, temperature and humidity fluctuations. Rajasthan is northwestern state of India, which has shown several dengue outbreaks in the past. In this paper we have tried to analyze the effects of annual cumulative rainfall on Dengue incidence in one of the largest and severely affected states of India. Retrospective data for Dengue incidence and Rainfall for the state of Rajasthan was collected and Pearson’s coefficient correlation was calculated as a measure of association between the variables. Our results indicate that annual cumulative rainfall shows a strong positive correlation with dengue incidence in the state of Rajasthan. Such analyses have the potential to inform public health official about the control and preparedness for vector control during monsoon season. This is the first study from the Indian state of Rajasthan to assess the impact of annual rainfall on dengue incidence, which has seen several dengue outbreaks in the past.

## Introduction

Dengue fever, an arboviral disease has become a significant public health problem in the last three decades. It has found new geographical niches, establishing itself as an endemic disease [1]. Nearly, 3.9 billion people in 128 countries are at risk of developing Dengue fever [2]. Substantial number of disease burden is contributed from India. In 2015 nearly 100,000 cases were reported from all the major Indian states and union territories [3]. The disease is caused by dengue virus of the family *Flaviviridae* and transmitted through an arthropod vector. The female of the *Aedes sp.* mosquito transmits the virus as it sucks the blood for nurturing the eggs. The infection might remain asymptomatic or develop in dengue fever (DF) or a more severe form of Dengue hemorrhagic fever (DHF) [4]. In absence of any potent anti-virals, disease management and prevention is dependent upon supportive therapy and control of vector population [5,6]. The disease seems to have seasonal pattern, with monsoon season witnessing a large number of cases. The Indian sate of Rajasthan is an arid desert, which receives rainfall during the monsoon period starting from July to September. It is in this season that most of the dengue cases are reported. First documented study of Dengue in Rajasthan was recorded as far back as 1969 [7] and since then many cases of Dengue have been reported. Several studies have identified weekly and monthly rainfall patterns, temperature and humidity fluctuations as important factors contributing to the survival and dispersal of vector population [8-10]. Such studies have the potential to alarm public health agencies for impending outbreaks. In this study we have analyzed the correlation between annual cumulative rainfall received and Dengue incidence for the Indian state of Rajasthan.

## Materials and method

The city of Rajasthan is located 26.57268^°^N and 73.83902^°^E in northwestern India. It is the largest state geographically covering an area of 342,239 km^2^. According to the 2011 census the population of Rajasthan is estimated to be 74,791,568 and population density is 200/km^2^. The climactic conditions of Rajasthan can be termed as dry and arid as vast swathes of land are covered with desert. Rainfall is received during the months between July to September. Winter, Pre-monsoon and Post-monsoon sees very little rainfall.

The data for dengue incidence for Rajasthan was obtained from National vector borne disease control program (NVBDCP). Retrospective data of the past ten years from 2005 to 2015 was collected. Climate data for Rajasthan was obtained by the yearly weather reports published by Indian Metrological Department (IMD). Total annual rainfall along with winter, Pre-monsoon, monsoon and post-monsoon rainfall data were considered for the final analysis. Pearson coefficient was calculated to identify correlation between dengue cases and rainfall. Rainfall was considered as independent variables while annual number of dengue cases was considered as dependent variable.

## Results

Retrospective data obtained from NVBDCP for the last 11 years (2005-2015) indicated that a total of 17008 cases and 108 deaths due to Dengue were reported from Rajasthan (Table 1). Maximum number of cases (n=4413) was reported in the year 2013 and minimum number of cases (n=370) was reported in the year 2005 (Figure 1). The year 2006 witnessed maximum number of deaths (n=26) due to Dengue. Most parts of Rajasthan are an arid desert with hot and sub-humid climate. Rainfall is sparse and takes place in the months of July, August and September. An average of 92 percent rainfall took place in the monsoon season (Figure 2) with the year 2011 reporting maximum rainfall (638 mm) and the year 2009 reporting minimum rainfall (314 mm). An analysis of correlation between annual cumulative rainfalls with incidence of dengue was carried out by calculating Pearson’s correlation coefficient (Table 2). An overall positive and moderately strong correlation (R=0.51) was found between the two variables indicating that rainfall does affect annual cases of Dengue in Rajasthan. The monsoon season receives maximum rainfall, and a concomitant rise in dengue cases has been observed. Analysis of dengue incidence with rainfall received during different seasons showed that the strongest correlation between dengue incidences was with rainfall pattern during monsoon season (Figure 1).

**Table 1.** Number of annually reported cases of Dengue, deaths and annual rainfall for the state of Rajasthan.

| Year | Dengue cases | Deaths | Rainfall |
| --- | --- | --- | --- |
| 2005 | 370 | 5 | 438.8 |
| 2006 | 1805 | 26 | 537 |
| 2007 | 540 | 10 | 417.9 |
| 2008 | 682 | 4 | 468 |
| 2009 | 1389 | 18 | 313.6 |
| 2010 | 1823 | 9 | 607.35 |
| 2011 | 1072 | 4 | 637.85 |
| 2012 | 1295 | 10 | 507 |
| 2013 | 4413 | 10 | 612.15 |
| 2014 | 1243 | 7 | 492.7 |
| 2015 | 2376 | 5 | 554 |

**Figure 1.**
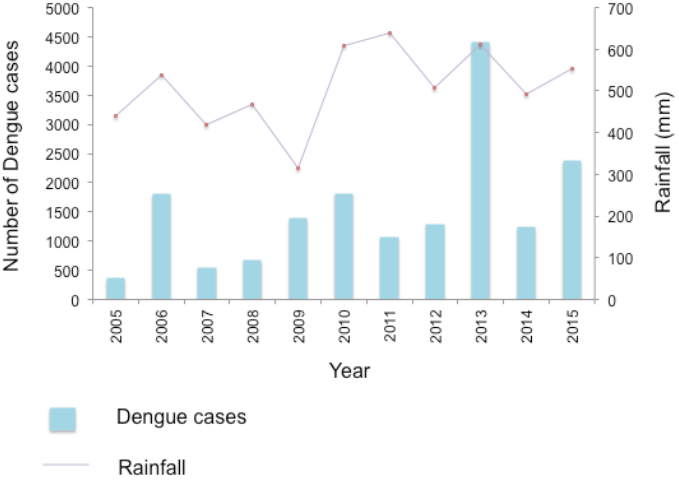
Comparison of dengue cases with annual cumulative rainfall

**Figure 2.**
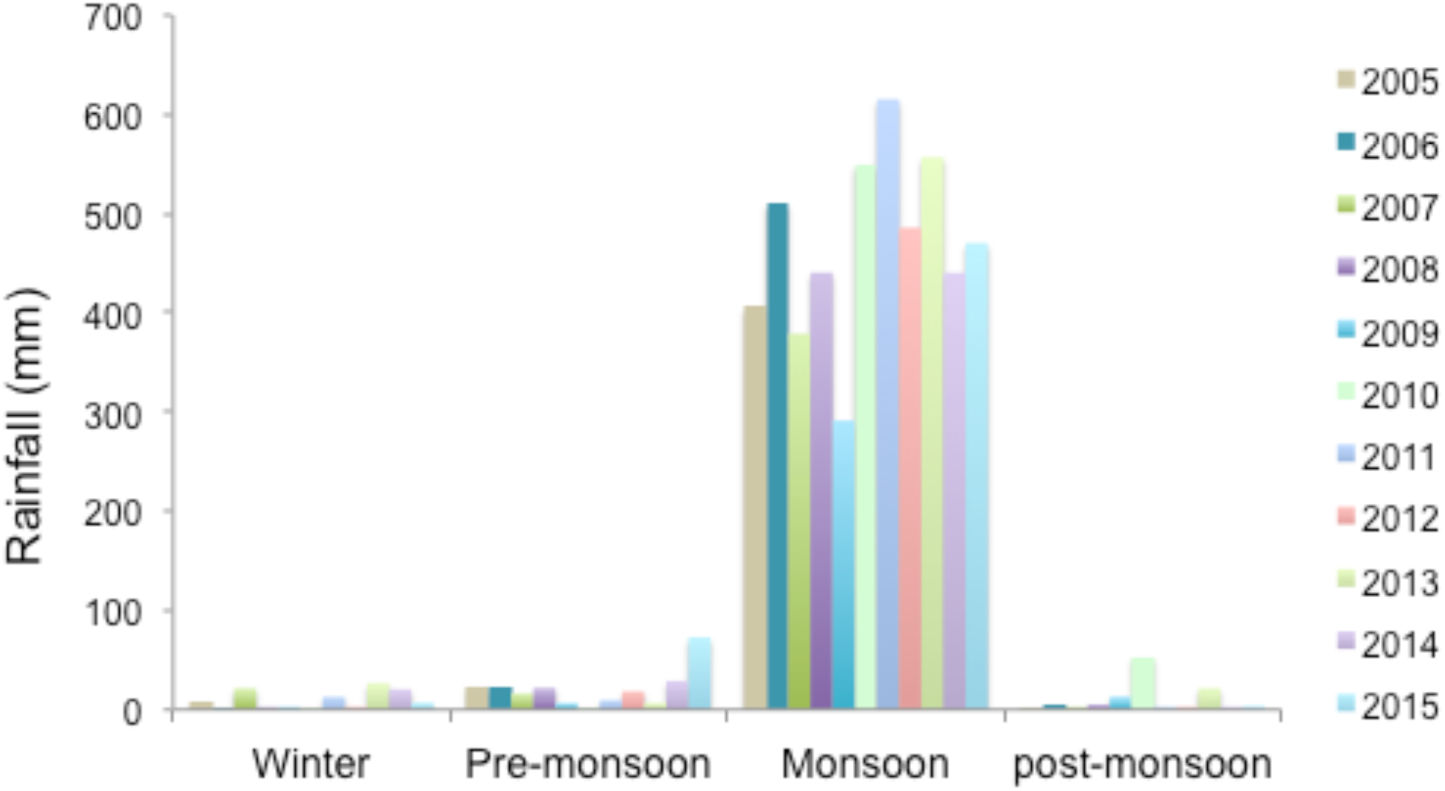
Season-wise distribution of annual cumulative rainfall

**Table 2.** Correlation coefficient between dengue incidence and total, winter, pre-monsoon, monsoon and post-monsoon rainfall

|  |  |  |
| --- | --- | --- |
|  |  | **Rajasthan** |
| **Rajasthan** | **R** | **1.** |
|  | *R Standard Error* |  |
|  | *t* |  |
|  | *p-value* |  |
| **Total Rainfall** | **R** | **0.51** |
|  | *R Standard Error* | 0.082 |
|  | *t* | 1.779 |
|  | *p-value* | 0.109 |
| **Winter rainfall** | **R** | **0.352** |
|  | *R Standard Error* | 0.097 |
|  | *t* | 1.129 |
|  | *p-value* | 0.288 |
| **Pre-monsoon rainfall** | **R** | **0.015** |
|  | *R Standard Error* | 0.111 |
|  | *t* | 0.046 |
|  | *p-value* | 0.965 |
| **Monsoon rainfall** | **R** | **0.429** |
|  | *R Standard Error* | 0.091 |
|  | *t* | 1.426 |
|  | *p-value* | 0.188 |
| **Post-monsoon rainfall** | **R** | **0.405** |
|  | *R Standard Error* | 0.093 |
|  | *t* | 1.328 |
|  | *p-value* | 0.217 |

## Discussion

Dengue continues to be a public health threat in India. It is found to be endemic in all the states and union territories of India. Rajasthan is a northeastern state with the largest landmass most of which is covered in desert. The climate is characterized by little rainfall, high wind velocity and hyperthermic conditions. With scarcity of water, people have a tendency of storing water for domestic use. The earliest published reports of dengue from the state were reported in the year 1969. Since then the city has seen many outbreaks of Dengue in the last couple of years [11,12]. Gathering and availability of publically available data continues to be a stumbling block for the analysis of vector borne diseases in India. Though NVBDCP reports annual dengue incidence from each state separately, but lack of weekly or monthly breakup is still crucial in determining disease endemicity and reasons for sudden outbreaks. A major factor that contributes in disease incidence is rainfall. Over the years many reports have been published that indicate a seasonal pattern of dengue. In India most of the cases are reported during monsoon season, probably by providing optimal conditions for vector survivability. Weekly rainfall, temperature and humidity fluctuation have been shown to effect disease incidence however [13-17], there are conflicting reports on the affects of annual rainfall on Dengue incidence. We have analyzed the correlation between annual rainfall as reported by IMD in their yearly-published reports and annual dengue incidence for the state of Rajasthan. Our analysis indicates a moderate positive correlation between annual rainfall and dengue cases. We also analyzed correlation between rainfall received during winter, pre-monsoon, monsoon and post-monsoon season. Our results indicate an overall positive and strong correlation between dengue incidence and rainfall received during monsoon season. These results may not indicate causality but do suggest strong positive correlation between monsoon rainfall and dengue incidence. Evidently, climactic factors in the state of Rajasthan influence dengue incidence. Our study, though preliminary in nature, does point out annual rainfall as an important predictor of dengue incidence in Rajasthan. We do acknowledge the limitations of our study as weekly rainfall and dengue patterns are not available from either IMD or NVBDCP, which are better variables to be considered for analysis. Nonetheless, in the dry and arid regions of Rajasthan where water availability is scarce and for most part of the year weather is generally very hot, monsoon season rainfall is a major contributor towards incidence of dengue. In absence of any therapeutic intervention, the control of Dengue is heavily dependent upon the control of vector population. Knowledge of weather variables that might influence disease incidence would be helpful in the control and limitation of Dengue incidence.

## Conclusion

Weather variables are important determining factors for arboviral diseases. Prevalence of dengue during monsoon season in many parts of the world indicates a climatic correlation. Present study is an attempt at understanding the affects of annual rainfall on dengue incidence. By calculating Pearson’s correlation coefficient, we show a moderately strong positive influence of rainfall on dengue. Though preliminary in nature, our data justifies a need for a more thorough analysis of weather variables on dengue incidence. Additionally, public availability of weekly and monthly surveillance data for weather variables and Dengue incidence from Rajasthan and other parts of country will be helpful in predicting dengue outbreaks. Such analysis will help public health agencies in designing better approaches for the control of Dengue.

## Acknowledgements

N/A.

## Availability of data and materials

The datasets analysed during the current study is available from the corresponding author on reasonable request.

## Authors’ contributions

NS collected and analyzed the data and wrote the manuscript

## Consent for publication

N/A.

## Ethics approval and consent to participate

/A.

